# Membrane integration and topology of RIFIN and STEVOR proteins of the *Plasmodium falciparum* parasite

**DOI:** 10.1101/653998

**Authors:** Annika Andersson, Renuka Kudva, Anastasia Magoulopoulou, Quentin Lejarre, Patricia Lara, Peibo Xu, Suchi Goel, Jennifer Pissi, Xing Ru, Tara Hessa, Mats Wahlgren, Gunnar von Heijne, IngMarie Nilsson, Åsa Tellgren-Roth

**Affiliations:** Department of Biochemistry and Biophysics, Stockholm University, SE-10691 Stockholm, Sweden; Center for Infectious Disease Research, Department of Microbiology, Tumor and Cell Biology, Karolinska Institutet, Stockholm, Sweden; Science for Life Laboratory, Stockholm University, Box 1031, SE-171 21 Solna, Sweden

**Keywords:** RIFIN protein, STEVOR protein, oligosaccharyltransferase complex, protein synthesis, transmembrane domain, membrane protein, membrane topology, membrane protein biogenesis, parasite, red blood cell

## Abstract

The malarial parasite *Plasmodium*, infects red blood cells by remodeling them and transporting its own proteins to their cell surface. These proteins trigger adhesion of infected cells to uninfected cells (rosetting), and to the vascular endothelium, obstructing blood flow and contributing to pathogenesis. RIFINs (*P. falciparum*-encoded repetitive interspersed families of polypeptides) and STEVORs (subtelomeric variable open reading frame), are two classes of proteins that are involved in rosetting. Here we study the membrane insertion and topology of three RIFIN and two STEVOR proteins, employing a well-established assay that uses N-linked glycosylation of sites within the protein as a measure to assess the topology a protein adopts when inserted into the ER membrane. Our results indicate that all the proteins tested assume an overall topology of N_cyt_-C_cyt_, with predicted transmembrane helices TM1 and TM3 integrated into the ER membrane. We also show that the segments predicted as TM2 do not reside in the membrane. Our conclusions are consistent with other recent topology studies on RIFIN and STEVOR proteins.

---

Malaria is a major disease plaguing the tropical and subtropical parts of the world, with infants, children, pregnant women, and immune-compromised adults being the most vulnerable to infection (1). The most fatal form of malaria in humans, caused by *Plasmodium falciparum*, is characterized by obstructed blood flow in the microvasculature that results from the adhesion of infected red blood cells (iRBC) to the vascular endothelium, and rosetting of iRBCs to healthy RBCs (2,3). The consequences include oxygen deprivation, organ failure, stillbirth in cases of placental malaria, and cerebral malaria. Rosetting and adhesion of iRBCs has been linked to the display of pathogen-derived variant surface antigens (VSAs) on their cell surface (reviewed in (4)). The most extensively studied VSA is PfEMP-1 (*P. falciparum* erythrocyte membrane protein 1), and others include the RIFIN (repetitive interspersed family) and STEVOR (subtelomeric variable open reading frame) proteins.

RIFINs are encoded by the rif-gene family, comprising around 150 genes (5), and belong to the two-transmembrane (2TM) superfamily (6), although several of them assume a one-transmembrane (TM) topology in the iRBC plasma membrane (6,7). Two groups of RIFINs, A and B, have been described so far. RIFIN-A proteins (70% of all RIFINs) are characterized by the presence of 25 amino acids at the N-terminus that are lacking in RIFIN-B proteins (8). The former are exported from the parasite to the surface of RBCs, while RIFIN-B proteins are mainly localized within the parasite and therefore potentially serve a different function (8,9). STEVOR proteins, expressed in the merozoite stage of the parasite’s life cycle (10), have been proposed to either have a single TM domain (7) or two TMs (5). They have been shown to mediate RBC rosetting independently of PfEMP1 (10).

RBCs lack cell organelles and vesicular trafficking pathways, making it essential for the parasite to remodel them completely to ensure its own survival. Immediately following invasion, the parasite is enclosed in a parasitophorous vacuole (PV) between two membranes: the PV membrane (PVM) and the parasite plasma membrane (PPM) respectively. Thereafter, protein export occurs in steps; they are first synthesized and then secreted across the parasite membrane to the PV using the secretory pathway. The *P. falciparum* genome contains homologs of the Sec translocon in the endoplasmic reticulum (ER) membrane and other factors important for targeting and translocation of proteins (11). Nascent polypeptides can be modified by the cleavage of signal peptides by signal peptidase (12) or by N-linked glycosylation by the oligosaccharyl transferase (OST) enzyme complex (13), however the significance of the latter is unclear (14). A large portion of exported proteins are cleaved in the ER (15,16) by a *Plasmodium*-specific enzyme called Plasmepsin V (17) at an N-terminal motif (RxLxE/Q/D) called the PEXEL (*Plasmodium* export element) site (15). Further transport of proteins to the iRBC cytosol and surface occurs via pathways set up by the parasite (reviewed in (18,19)). It is therefore important to describe the biogenesis of parasite proteins due to their well-established connection to pathogenesis, disease severity and parasite survival.

A previous study from our lab on one RIFIN-A and RIFIN-B protein expressed *in vitro* with canine pancreas-derived rough ER membranes (microsomes), demonstrated that the C-terminal transmembrane TM helix is membrane-inserted with the N-terminus oriented towards the lumen and the C-terminus in the cytosol (N_lum_-C_cyt_) (20). The loop between the N-terminal hydrophobic domain (that is believed to be a signal peptide (SP)) and the C-terminal TM helix was shown to be located in the ER lumen (20). In this study, we further investigate the topology of the previously studied RIFIN proteins (20), an additional RIFIN-A, and two STEVOR proteins using a well-established *in vitro* method (21,22). Glycosylation-competent green fluorescent protein (gGFP)-tagged RIFIN and STEVOR proteins were also visualized in a human cell line to complement the data obtained from the *in vitro* experiments. We show that these proteins all exhibit an overall N_cyt_-C_cyt_, topology in the ER membrane.

## Results

### Prediction of TMs of RIFIN and STEVOR proteins

We used the ΔG predictor (http://dgpred.cbr.su.se) (26) to predict potential TM domains for all the proteins studied. RIFIN-A1 (RA1, 372 residues-long) was predicted to have two putative TM (pTM) domains between residues 1-19 and 331-353, RIFIN-A2 (RA2, 330 amino acids) was predicted to have three pTM domains between residues 1-18, 137-159 and 291-313, RIFIN-B (RB, 338 amino acids), three pTM domains between residues 3-25, 119-141, and 297-319. STEVOR 06 (S06, 306 residues long) was predicted to have three pTM domains between residues 3-21, 179-201 and 265-287, and STEVOR 10 (S10, 305 amino acids in length) three pTM domains between 3-21, 179-201 and 268-290.

### Membrane integration efficiency of RIFIN proteins

#### Experimental setup

The RIFIN and STEVOR proteins were all translated *in vitro*, in the presence and absence of canine ER-derived column-washed rough microsomes (CRM) to probe for membrane insertion. N-linked glycosylation of specific sites within the protein sequence (NXS/T) was used to distinguish between luminal and cytosolic domains of the protein. N-glycosylation is catalyzed by the OST that is active on the luminal side of the ER membrane (13). Glycosylated products migrate slowly on SDS-PAGE compared to unmodified proteins, with one glycan adding to ~2.5 kDa to the molecular mass of proteins. Glycosylation of proteins was confirmed by digestion by Endo H.

We also engineered the RIFIN and STEVOR proteins or derivatives thereof into our well-established experimental “host” proteins derived from the *E.coli* LepB protein (25) – LepH2 and LepH3- and assessed the topology of inserted pTMs by N-linked glycosylation (21,22). In brief, LepB contains two natural TM domains (H1 and H2) and a C-terminal globular domain (P2). H2 or H3 were substituted by our protein sequence of interest together with a spacer of three glycines and one proline residue (GPGG) to shield the insert from the model protein. The H2 model protein allows the analysis of membrane insertion efficiency of the integrated domain with its N-terminus in the cytosol and C-terminus in the lumen (N_cyt_-C_lum_) (22). In the H3 model protein, the engineered peptide of interest will assume an N_lum_-C_cyt_ orientation in the membrane (21,26). Unidentified protein products are indicated by an (*) on the gel images.

#### Topologies of full-length RIFIN and STEVOR proteins

The topologies of full-length wildtype (WT) RA1 and RB have been previously examined (20). In the present study, we evaluated the topology of full-length RA2 (1-330) *in vitro*. RA2 has two naturally occurring potential glycosylation acceptor sites (GSs) in its sequence at positions 232 and 233 i.e. in the loop between pTMs 2 and 3. Since these GSs are adjacent to one another, they are potentially too closely spaced to be glycosylated at the same time (supplementary Figure S1) (32). The cytosolic or luminal locations of the other two loops were determined by engineering additional GSs at position 61 (N_61_ST) between pTM1 and pTM2, and at position 326 (N_326_TT) at the C-terminus of pTM3 (Figure 1A) respectively. WT RA2 translated in the presence of CRMs was mono-glycosylated and RA2 with an engineered GS i.e. N_61_ST was di-glycosylated, (Figure 1B, left panel). RA2 with the both the natural and two artificial GSs N_326_ and N_61_ respectively was also di-glycosylated (Figure 1B, right panel), indicating that only two sites are occupied by glycans and not the third one. These results suggest that the C-terminus of RA2 is located in the cytosol, while the loop between pTM1 and pTM3 is in the lumen, indicating that pTM3 inserts into the membrane in an N_lum_-C_cyt_ orientation and that pTM2 is not a transmembrane domain.

**Figure 1.**
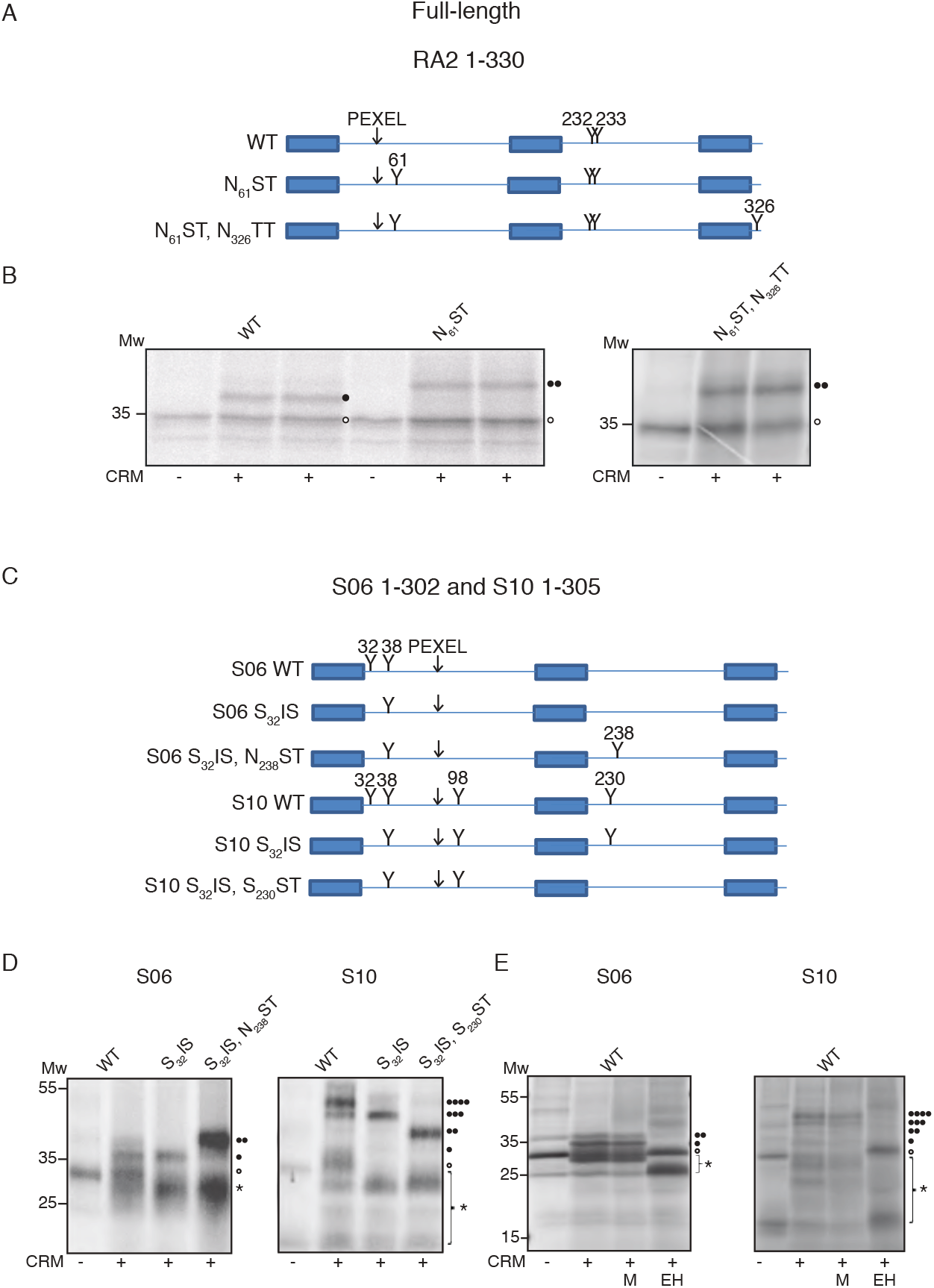
Investigating the topology of full length RIFIN and STEVOR proteins in the membrane: (A) Schematic of full-length RIFIN-A2 (RA2) with the PEXEL site and positions of glycosylation sites indicated by a (Y) and the residue number. (B) *In vitro* translated products of WT full-length RA2 and a mutant with disrupted GSs. Translation was carried out in the absence and presence of column-washed rough microsomes derived from canine pancreas. (○) represents unglycosylated protein species and glycosylated bands are represented by (•). (C) Schematic representations of STEVOR (S06 and S10) proteins with PEXEL and glycosylations sites indicated. (D) *In vitro* translated products of WT full-length S06 and S10, and mutants with disrupted GSs. Translations were carried out in the absence and presence of CRMs as indicated by (−/+). (○) represents unglycosylated protein species, all glycosylated products indicated by multiple (•), and protein bands of unknown identity indicated by (*). (E) *In vitro* translations of S06 and S10 in the absence and presence of CRMs. +CRM samples were treated with EndoH (EH) to digest glycosylated products. (M) represents the mock where samples were treated exactly the same as EH, but were not supplemented with the enzyme.

Full-length WT S06 has two natural GSs between pTM1 and pTM2, at positions N_32_ and N_38_ (Figure 1C). WT S06 translated in the presence of microsomes yielded proteins corresponding in molecular mass to unmodified S06, and mono-glycosylated and weakly di-glycosylated adducts, the latter of which we believe results from the GS at N_32_ being inefficiently glycosylated (Figure 1D, left panel). Perturbing this GS by mutating N_32_ to a serine (construct S_32_IS), resulted in a mono-glycosylated protein, allowing us to conclude that the loop between pTM1 and pTM2 is located in the lumen of the ER. Introducing a GS between pTM2 and pTM3 in construct S_32_IS, at position 238 (N_238_ST) to investigate where the loop between them is located, resulted in a di-glycosylated protein (Figure 1D, left panel) indicating that the loop between pTM2 and pTM3 is also located in the lumen.

WT full-length S10 has four natural GSs in total (Figure 1C), three sites located between pTM1 and pTM2 at positions N_32_, N_38_ and N_98_ and one between pTM2 and pTM3 at position N_230_. WT S10 translated in the presence of microsomes was tri- and tetra-glycosylated (Figure 1D, right panel). Perturbing the GS at position 32 (i.e. construct S_32_IS) yielded a tri-glycosylated species, and removing the GS at N_230_ (construct S_230_ST) in order to locate the loop between pTM2 and pTM3 resulted in a di-glycosylated protein. Glycosylation of proteins was confirmed by digestion by Endo H (Figure 1E).

Our results for RIFIN-A2 and STEVORs S06 and S10 demonstrate that the loops between pTM1 and pTM2, and between pTM2 and pTM3, respectively, are localized in the lumen. This suggests that pTM2 is likely not a TM domain because it must be translocated to the lumen to accommodate localization of the loops.

#### Expression of full-length RIFIN and STEVOR proteins engineered into the model protein LepH2

In order to further map the orientation of RIFIN and STEVOR pTM1 and pTM3 in the membrane, we engineered these proteins into our model LepB system. We chose to engineer full-length RIFINs (RA1, RA2 and RB) and STEVORs (S06 and S10) at position H2 in the LepB model protein (Figure 2A), based on our previous observations that these proteins most likely insert in an N_cyt_-C_cyt_ orientation in the membrane. Our current analysis was expanded to include the previously studied RA1 and RB proteins (20).

**Figure 2.**
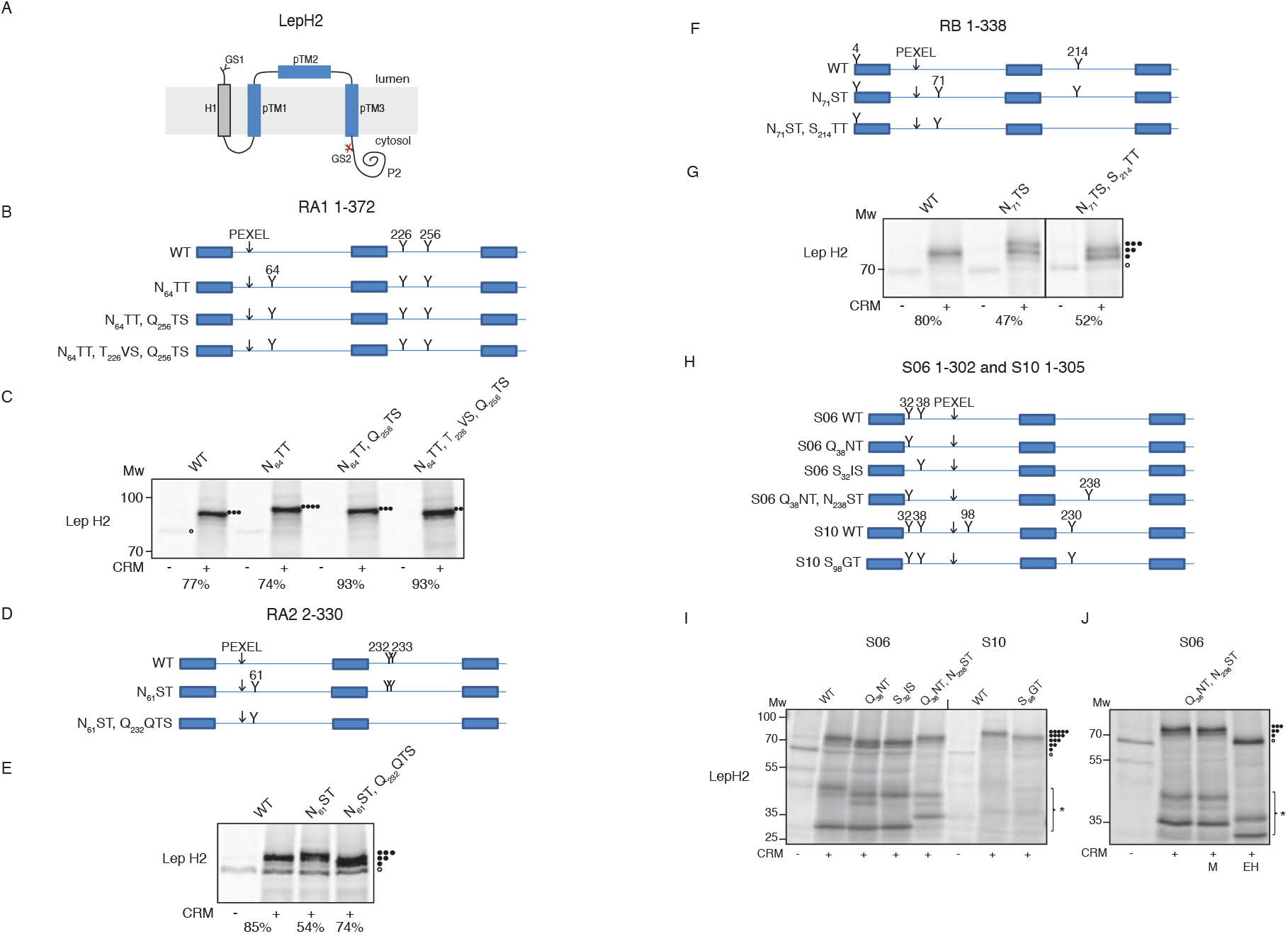
Investigating the topology of full length RIFIN and STEVOR proteins when engineered into LepH2. (A) Schematic representation of WT RA and STEVOR proteins engineered into LepH2. TM1 of LepB indicated as H1, and putative TM domains of RIFIN and STEVOR proteins indicated as pTM 1, 2 and 3. Glycosylation sites (GS) indicated by (Y). (B) Schematic of full-length RA1 and mutants with PEXEL and glycosylation sites indicated. (C) *In vitro* translated products of WT and mutant RA1 engineered at LepH2. All glycosylated products indicated by multiple (•). (D) Schematic of full-length RA2 and mutants with PEXEL and glycosylation sites indicated. (E) *In vitro* translated products of WT and mutant RA2 engineered at LepH2. All glycosylated products indicated by multiple (•). (F) Schematic of full-length RA1 and mutants with PEXEL and glycosylation sites indicated. (G) In vitro translated products of WT and mutant RB engineered at LepH2. All glycosylated products indicated by multiple (•) (H) Schematic of full-length S06 and S10 and their respective mutants with PEXEL and glycosylation sites indicated. (I) *In vitro* translated products of WT and mutant S06 and S10 engineered at LepH2. All glycosylated products indicated by multiple (•). (J) *In vitro* translations of a mutant of S06 translated in the absence and presence of CRMs. +CRM samples were treated with EndoH (EH) to digest glycosylated products. (M) represents the mock where samples were treated exactly the same as EH, but were not supplemented with the enzyme.

The two naturally occurring GSs in RA1 (N_226_VS and N_256_TS) and the two GSs in the LepH2 model proteins result in a total of four possible sites that can be glycosylated (Figure 2A and B). *In vitro* expression of RA1-H2 resulted in a 77% tri-glycosylated product (Figure 2C). An engineered GS at position 64 (N_64_TT) between pTM1 and pTM2 resulted in a tetra-glycosylated protein (74%) (Figure 2C). Individually disrupting the two natural GSs between pTM2 and 3 in construct N_64_TT to T226VS and Q256TS, resulted in equal proportions of tri-glycosylated and di-glycosylated proteins (Figure 2C). We conclude that the predominant topology of RA1-H2 is where both pTM1 and pTM3 span the membrane, and pTM2 is in the lumen (Figure 2A), consistent with our findings for full-length RA2 shown above. We observed that RA2-H2 also inserted with the same topology as RA1-H2, also confirming our previous observations (Figure 2D and E).

The two natural GSs in RB (N_4_ and N_214_TT) and those in LepH2 give a total of four potential glycosylation sites (Figure 2F). Translation of RB-H2 in the presence of CRMs resulted in predominantly di-glycosylated (80%) species (Figure 2G), indicating four conceivable topologies. An engineered GS (N_71_TS) between pTM1 and pTM2, resulted in an additional tri-glycosylated adduct (47%). Thus, in about half the translated protein, the loop between pTM1 and pTM2 is translocated into the lumen. A S_214_TT mutation resulted in a di-glycosylated protein (52%), leaving us with two probable conformations, one where pTM3 is membrane-integrated in its preferred orientation, and the other with either pTM1 or pTM2 integrated (Figure 2G).

The WT S06-H2 construct has a total of four GSs (including the GSs within LepH2 and S06), indicated in Figure 2H. Translating S06-H2 resulted in a tri-glycosylated protein (Figure 2I) and disrupting the GS at N_38_ (construct Q_38_NT), yielded a di-glycosylated protein instead, similar to what was seen for the S_32_IS construct (Figure 2I) where N_32_IS is disrupted. This confirms that both N-terminal GSs of S06 are glycosylated, although the modification of N_32_IS is inefficient. An additional GS at position 238 (N_238_ST) resulted in a tri-glycosylated product, indicating that pTM1 and pTM3 are integrated into the ER membrane, and pTM2 resides in the ER lumen. WT S10-H2 is penta-glycosylated, at four GSs in S10 and one in LepH2 (Figure 2H, I). A mutant of S10-H2 where the GS at N_98_ (construct S_98_GT) is disrupted was tetra-glycosylated (Figure 2I), indicating that the loop between pTM1 and 2 is located in the lumen. Glycosylation of proteins was confirmed by digestion by Endo H (Figure 2J).

These experiments demonstrate that RA1, RA2, and S06 insert in an N_cyt_-C_cyt_ orientation in the membrane, but do not indicate the orientation the pTM domains of RB and S10 can adopt when inserted. We therefore chose to also engineer single pTM regions of all the proteins into LepH2 and LepH3 to allow them to insert in their preferred orientations in the membrane.

#### Integration of single pTMs of RIFIN and STEVOR proteins when engineered into the LepH2 and LepH3 model proteins

To further validate the above results, we engineered single pTMs 1 and 3 of all three RIFIN proteins and the two STEVOR proteins with their flanking regions into the model proteins, LepH2 and LepH3 (Figure 3A).

**Figure 3.**
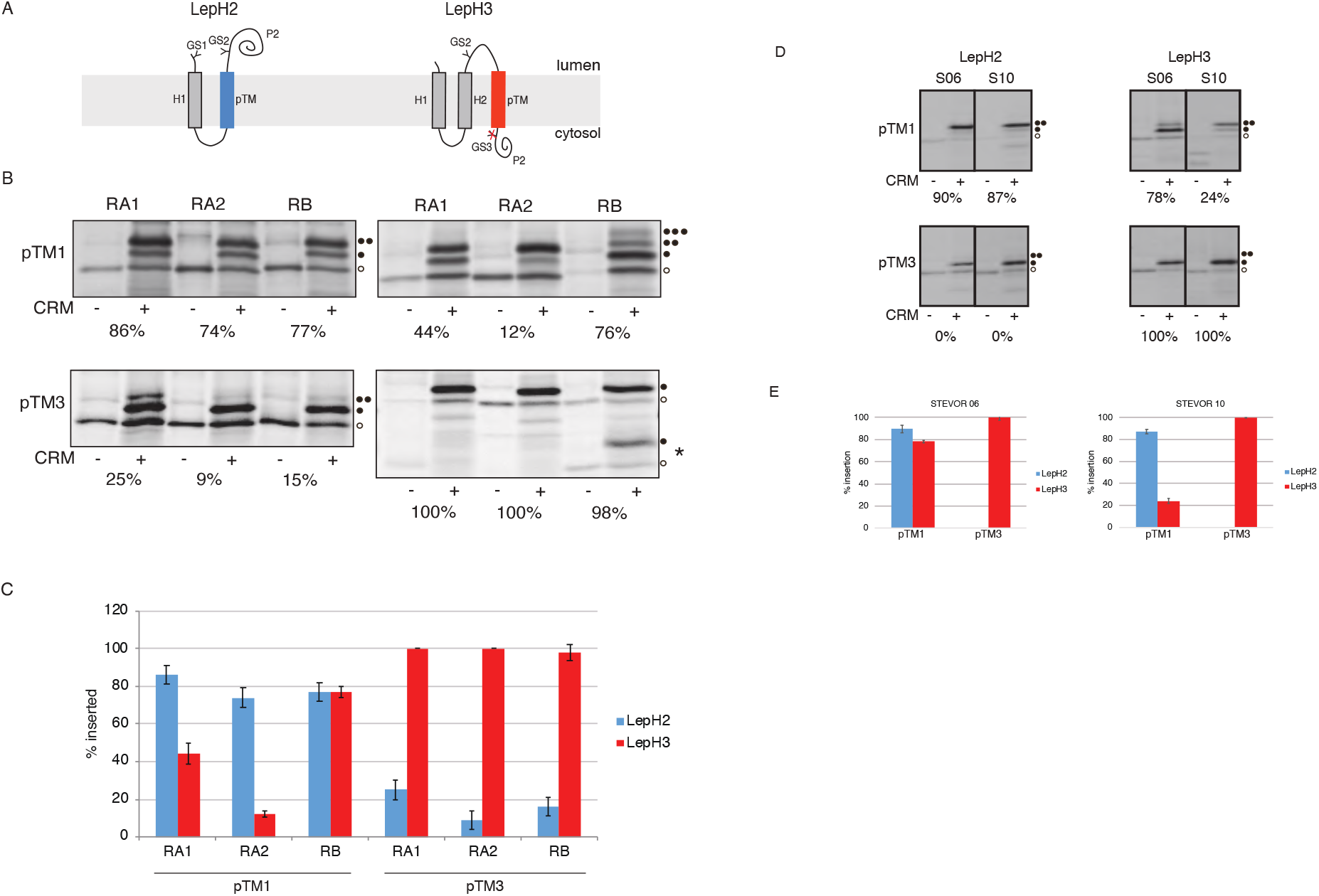
Insertion and topology of single pTMs of RIFIN and STEVOR proteins when engineered into LepH2 and LepH3. (A) Schematic representing single TMs of RIFIN and STEVOR proteins when engineered into LepH2 (left panel) and LepH3 (right panel). (B) *In vitro* translations of single pTMs of RA1, RA2 and RB engineered into LepH2 (left panels) and LepH3 (right panels), glycosylated products represented by (•). (C) Graphical representations of the insertion efficiencies of single pTMs of RA1, RA2 and RB as assessed by glycosylation. Error bars represent standard deviations calculated from an average of three independent experiments. (D) *In vitro* translations of single pTMs of S06 and S10 engineered into LepH2 (left panels) and LepH3 (right panels), glycosylated products represented by (•), and protein bands of unknown identity indicated by (*) (E) Graphical representations of the insertion efficiencies of single pTMs of S06 and S10 as assessed by glycosylation. Error bars represent standard deviations calculated from an average of three independent experiments.

pTM1 of both RA1 and RA2 when engineered in the H2 position integrated efficiently (86% and 74% respectively) in an N_cyt_-C_lum_ topology in the ER membrane, (Figure 3B, left panel, Figure 3C). Conversely, the same pTMs were poorly integrated (44% and 12% respectively) when engineered in the H3 position (Figure 3B, right panel, Figure 3C). pTM1 of RB was well-inserted into the membrane both in the H2 and H3 positions (approx. 77%), i.e. it can adopt two possible orientations in the membrane (Figure 3B). We also observed that a small proportion of RB pTM1 engineered at position H3 was translocated to the lumen, as indicated by the presence of a tri-glycosylated product, due to glycosylation of GS at N_4_ (Figure 3B, right panel).

pTM3 of all three RIFIN proteins integrated efficiently (to approx. 100%) in an N_lum_-C_cyt_ topology when engineered in the H3 position (Figure 3B, right panel, Figure 3C), and poorly when engineered in the H2 position (RA1 25%, RA2 9% and RB 15%, Figure 3B, left panel, Figure 3C), indicating that the preferred topology for pTM3 is N_lum_-C_cyt_, further supporting that full-length RA1, RA2 and RB insert in an N_cyt_-C_cyt_ topology in the membrane.

pTM1 of S06 were efficiently inserted in the membrane in both orientations (Figure 3D and E). pTM1 of S10 on the other hand integrated efficiently into the membrane when engineered in the H2 position (87 %), but not in the H3 position (24 %) (Figure 3D and E). Thus, S10 pTM preferentially orients N_cyt_-C_lum_, but S06 pTM1 does not exhibit any orientation preference.

pTM3 of both S06 and S10 integrate in an N_lum_-C_cyt_ orientation, i.e. when inserted in the H3 position, but do not integrate when engineered in the H2 position (Figure 3D and E), consistent with previous observations that both flanking charges and hydrophobicity determine the orientation of a TM fragment in the membrane (33–38), and with the finding that the overall topology of full-length STEVOR proteins is N_cyt_-C_cyt_.

#### Investigation of the topology of RIFINs and STEVORs in HEK293T cells

We finally wanted to verify whether the topologies that were established with our *in vitro* assay could be confirmed in human cells, using a previously described experimental set-up based on a glycosylation-competent green fluorescent protein variant (gGFP) (28). Briefly, each of the full-length RIFIN and STEVOR proteins was fused to gGFP at their C-terminus (Figure 4). gGFP fluoresces in the cytoplasm, but becomes glycosylated on two sites (preventing the maturation of the fluorophore) when translocated to the ER lumen, allowing us to track where the C-terminus of the protein of interest is localized. The RIFIN-gGFP and STEVOR-gGFP fusion constructs (RA1-gGFP, RA2-gGFP, RB-gGFP, S06-gGFP, and S10-gGFP) were transfected into HEK-293T cells and imaged 24 hours post-transfection (Figure 4B). As a control, we used SP-C-gGFP where gGFP translocates to the lumen and hence is non-fluorescent (28) (Figure 4C, right panel). We also visualized the expressed proteins in cell lysates by SDS-PAGE and Western blotting (Figure 4A).

**Figure 4.**
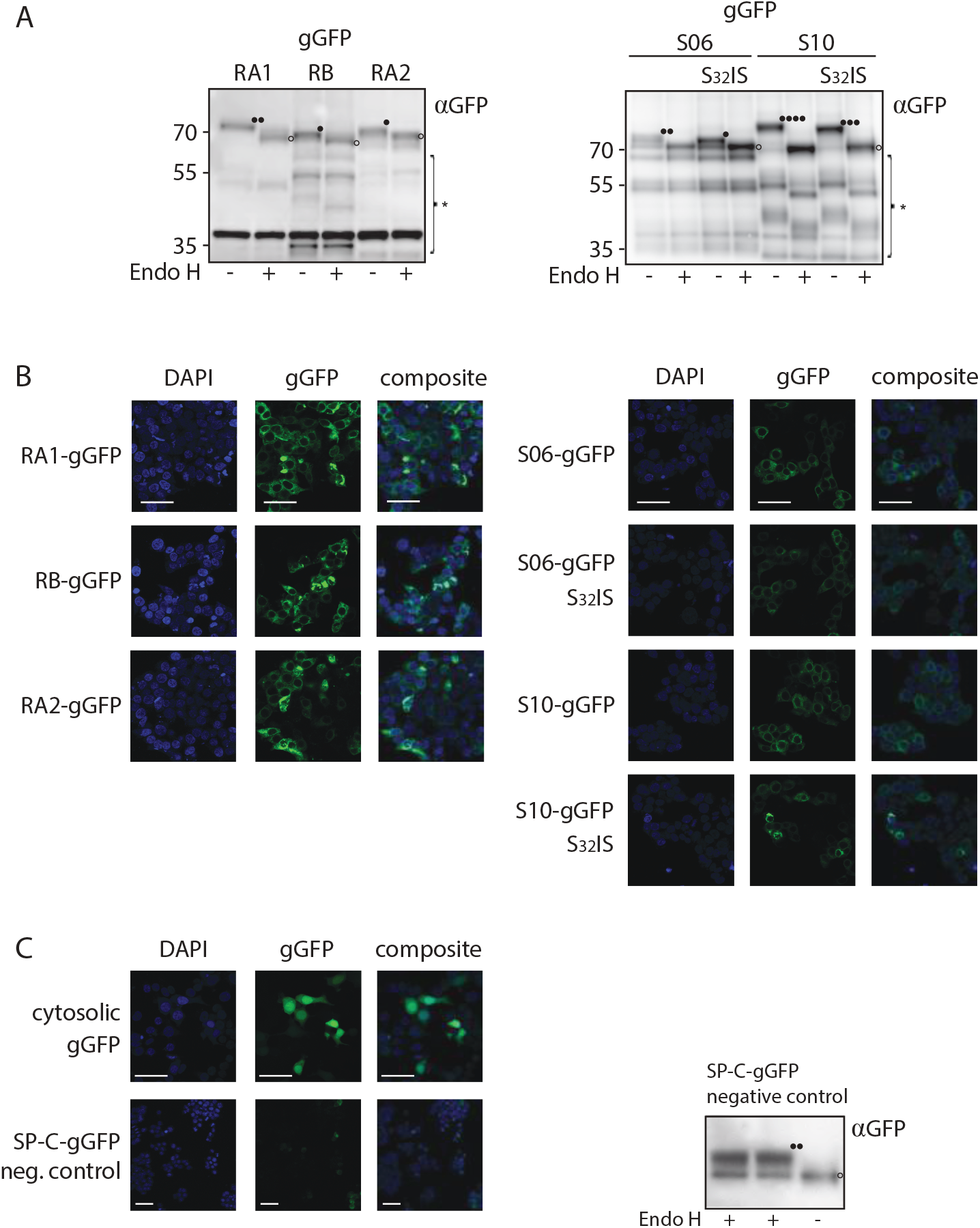
Visualizing topologies of RIFIN and STEVOR proteins in mammalian cells. (A) Glycosylation of RA1, RA2 and RB (left panel) and S06 and S10 (right panel) in cell lysates derived from HEK293T cells, assessed by Western blotting, glycosylated products represented by (•), and protein bands of unknown identity indicated by (*) Samples were treated with EndoH as a control. (B) The fluorescence pattern was examined by microscopy for both RIFIN (left panel) and STEVOR proteins (right panel), for both WT and mutants. (C) A positive (only gGFP) and negative control (SPC-gGFP) for the fluorescence (left panel), and the corresponding Western blot on the cell lysate for SPC-gGFP. DAPI staining, GFP and an overlay (composite) are indicated for the fluorescence images (Scale bars: 50 μm).

RA1-gGFP and RA2-gGFP fusions have a total of four GSs each, but due to the close proximity of the native GSs in RA2 (position 232 and 233) only three are potentially glycosylated. The RB-gGFP fusion protein has a total of four possible GSs, and three of them can be potentially glycosylated because one of them (N_4_) is located too close to the membrane to be accessible to OST. The fluorescence and glycosylation patterns observed (Figure 4A and B) indicate that the GSs within all the RIFIN proteins were translocated to the ER lumen, whereas the C-terminal gGFP remains in the cytosol and therefore is fluorescent (Figure 4C, left panel).

S06 has two native GSs at position 32 and 38, so the S06-gGFP protein has a total of four GSs. As a control, we expressed a protein where the GS N_32_IS in S06 was disrupted by the N32S mutation (S06-gGFP S_32_IS). The fluorescence (Figure 4B) and glycosylation pattern (Figure 4A) demonstrated that the GSs in S06 are in the ER lumen, and that gGFP lies in the cytosol. We also observed a similar pattern for S10-gGFP with its six GSs (four in S10 at position 32, 38, 98, and 230 and two in gGFP), where S10-gGFP S_32_IS served as a control.

These results confirm the findings from our *in vitro* experiments that pTM3 inserts into the membrane in an N_lum_-C_cyt_ orientation, and the mono-glycosylated species of pTM1 indicates that it is also inserted into the membrane.

## Discussion

In this study, we have determined the topologies of three RIFINs (RIFIN-A1 and 2, and RIFIN-B) and two STEVORs (STEVOR-06 and STEVOR-10) in a well-established *in vitro* assay supplemented with ER membranes. Although these membranes are derived from dog pancreas, the malarial parasite has a similar targeting and translocation system in its ER membrane (11,12) and so it is reasonable to assume that the two are comparable. However, the export of proteins in the parasite is a more complex process (39,40) because the proteins are secreted across several membranes to reach the surface of the iRBC, and most exported proteins are cleaved by the *Plasmodium* enzyme, plasmepsin V (15,41), which is absent in our system.

The ΔG predictor (26) predicts three potential TM domains for each of the proteins studied. We observe that pTM2, which is the least hydrophobic of the pTMs, does not reside in the membrane. pTM3, the most hydrophobic of the predicted TMs for all the proteins, appears to integrate the most efficiently in the membrane. pTM1 potentially serves as a signal sequence and targets RIFIN and STEVOR proteins to the ER membrane. Although our experimental system has a functional signal peptidase (22), it did not appear to cleave the proteins after pTM1. We cannot however rule out that cleavage takes place in the parasite (12).

We used our LepH2 model set-up to study how the full-length RIFIN and STEVOR proteins integrate into the membrane because it accommodated natural orientation of the proteins in the membrane, and this setup (21,22,26), allowed us to confirm that both pTM1 and pTM3 are membrane-integrated. The LepH2 and LepH3 systems also allowed us to probe the topology preferences of single pTMs in the membrane.

Our experiments indicate that pTMs1 of RA1, RA2, and S10 insert into the ER membrane, preferentially in an N_cyt_-C_lum_ orientation like a signal peptide. pTM1 of RB and S06, which are predicted to be the most hydrophobic of all potential pTM1s (Table 1), insert efficiently into the ER membrane and exhibit no orientation preferences, as shown by equivalent insertion efficiencies in both LepH2 and LepH3. pTM1 of S10 is slightly less hydrophobic, and its orientation in the membrane is potentially dictated by a lysine residue at the N-terminus, allowing it to adopt an N_cyt_-C_lum_ topology. In all five proteins studied, pTM3 was predicted to be strongly hydrophobic (Table 1), and we found that they integrated efficiently, with a strong tendency to adopt an N_lum_-C_cyt_ orientation, which is consistent with their hydrophobicity and charge distribution (all pTM3 segments include a number of positively charged residues at their C-terminal end). Our *in vitro* experiments, supported by the ΔG predictions, demonstrate that the studied RIFIN and STEVOR proteins all have two TM domains, pTM1 and pTM3.

**Table 1.** Position of the three pTM helices determined of each RIFIN and STEVOR by the ΔG predictor. Data shown represent predicted vs. measured ΔG values for each of the selected hydrophobic domains in RIFIN and STEVOR. For the predicted values the ΔG prediction server (http://www.cbr.su.se/DGpred/) was used. All pTMs were predicted from the full-length RIFIN and STEVOR protein and selected to be around 20-residues long using length corrections and sub-sequences with lowest ΔG. The number in the column measured positions correlate with the pTMs with flanking residues. All pTMs with flanks were insulated by a tetra-peptide, GGPG…GPGG. Integration (%) correlate with pTMs with flanking residues. The integration is calculated from at least three independent experiments.

| RIFIN | Region | Predicted position | Measured position | Predicted $\Delta G$ (kcal/mol) | H2 Integration (%) | H2 Measured $\Delta G$ (kcal/mol) | H3 Integration (%) | H3 Measured $\Delta G$ (kcal/mol) |
| --- | --- | --- | --- | --- | --- | --- | --- | --- |
| RA1 | pTM1 | 1-19 | 1-39 | 2.2 | 86 | -1.1 | 44 | 0.1 |
|  | pTM2 | - | 119-222 | - | 0 | 2.7 | 7 | 1.5 |
|  | pTM3 | 331-353 | 315-372 | -5.1 | 25 | 0.6 | 100 | -2.7 |
| RA2 | pTM1 | 1-18 | 2-35 | 2.5 | 74 | -0.6 | 12 | 1.2 |
|  | pTM2 | 137-159 | 117-179 | 1.8 | 63 | -0.3 | 16 | 1.0 |
|  | pTM3 | 291-313 | 273-330 | -4.6 | 9 | 1.3 | 100 | -2.7 |
| RB | pTM1 | 3-25 | 1-45 | 0.5 | 77 | -0.7 | 76 | -0.7 |
|  | pTM2 | 119-141 | 107-166 | 1.4 | 58 | -0.2 | 34 | 0.4 |
|  | pTM3 | 297-319 | 277-338 | -5.0 | 15 | 1.0 | 98 | -2.3 |
| S06 | pTM1 | 3-21 | 2-33 | -0.3 | 90 | -1.3 | 78 | -0.7 |
|  | pTM2 | 179-201 | 159-221 | 0.7 | 88 | -1.2 | 77 | -0.7 |
|  | pTM3 | 265-287 | 246-302 | -5.0 | 0 | 2.7 | 100 | -2.7 |
| S10 | pTM1 | 3-21 | 2-33 | 0.4 | 87 | -1.1 | 24 | 0.7 |
|  | pTM2 | 179-201 | 159-221 | 0.5 | 87 | -1.1 | 63 | -0.3 |
|  | pTM3 | 268-290 | 249-305 | -6.4 | 0 | 2.7 | 100 | -2.7 |

pTM segments were predicted from full length RIFIN and STEVOR with the $\Delta G$ predictor (Hessa, 2007). pTM length was set to be around 20 residues for flanking pTMs. $\Delta G$ values in kcal/mol were predicted with the $\Delta G$ predictor

We also probed the topology of the RIFIN and STEVOR proteins in mammalian cells by constructing fusions to a glycosylation-competent GFP variant. The results were consistent with the *in vitro* data. Both indicate that RA1, RA2, RB, S06, and S10 adopt an N_cyt_-C_cyt_ topology in the membrane, with pTM1 and pTM3 spanning the membrane and pTM2 residing in the ER lumen.

Our results reaffirm that hydrophobicity and charge distribution (33–38) dictate how TM domains insert into the membrane, and we believe that in the context of malarial proteins this can dictate localization of proteins. Future studies to obtain structures of these proteins and others of the class located on the surface of the infected red blood cell, are important for the understanding of the invasion of the host and avoidance of the immune system.

## Experimental procedures

### Enzymes and Chemicals

Unless otherwise stated, all chemicals were obtained from Sigma-Aldrich Merck (USA). Plasmid pGEM1 and TNT SP6 Quick Coupled Transcription/Translation System were purchased from Promega (USA). All enzymes were from ThermoFisher Scientific (USA) except for Phusion DNA Polymerase (New England Biolabs (USA)). Deoxynucleotides were from Agilent Technologies (USA) and oligonucleotides were from Eurofins MWG Operon (Germany). [^35^S]-methionine to label proteins in during translation was from PerkinElmer (USA). All other reagents of analytical grade and obtained from Merck Millipore (Germany).

### DNA manipulations

A codon optimized RIFIN-A2 (PfIT_060035900 CDS) and the PF3D7_0617600 (STEVOR 06, S06) and PF3D7_1040200 (STEVOR 10, S10) genes were engineered into the pGEM1 vector using a XbaI/SmaI site together with a preceding Kozak sequence as previously described (23–25). The other previously studied RIFINs were engineered into the pGEM1 vector as described in (20). Putative TMs for the selected RIFIN and STEVOR proteins were predicted by the ΔG predictor server (V 1.0) (https://www.dgpred.cbr.su.se) (26).

Selected sequences were amplified by PCR with appropriate primers containing GGPG-flanks and restriction sites for SpeI and KpnI enzymes and introduced into the DNA of the well-characterized model protein leader peptidase (LepB) from *Escherichia coli*. Two LepB model proteins (LepH2 and H3) were used as previously described (21,22). Sequences for S06 and S10 that were translated without the LepB background were codon optimized to facilitate easy cloning and expression, whereas those engineered into LepB were non-optimized. In the cDNA of RA1, thymine at position 263 was changed using QuikChange™ Site-Directed Mutagenesis protocol from Stratagene (La Jolla, CA, USA) into a cytosine in order to remove a SpeI cleavage site. Glycosylation sites (GS) were added or removed (NXS/T, where X ≠ a proline residue) by site-directed mutagenesis. In most cases a Thr (T) was used as the hydroxy amino acid in the glycosylation sequon. All constructs were confirmed by sequencing of plasmid DNA at Eurofins MWG Operon (Germany).

For WT RB proteins with the C-terminal sequence, multiple protein products were observed on SDS-PAGE, so a putative ribosome stalling sequence of R_322_KKKM encoded by a polyadenylate (polyA) stretch (AGAAAAAAAAAAATG) at the C-terminus of the protein was changed to a non-polyadenylate stretch (27), in a selection of constructs for comparison.

The constructs expressed in mammalian cells with the gGFP fusion were cloned into pcDNA3.1 (Invitrogen) as previously described (28).

### *In vitro* transcription and translation

Constructs engineered into the pGEM1 vector with and without LepB were translated in the TNT SP6 Quick Coupled Transcription/Translation system in a total reaction volume of 10 μl (5 μl reticulocyte lysate, 150 ng of DNA template, 0.5 μl of [^35^S]-methionine (5 μCi), and 1 μl of column-washed canine pancreas-derived rough microsomes (CRM) (tRNA Probes, USA) (29) for 90 min at 30°C (30). Translated proteins were analyzed by SDS-PAGE and autoradiography. The protein bands were quantified using Image Gauge V 4.23 to obtain an intensity cross section that was fitted to a Gaussian distribution using EasyQuant (Rickard Hedman, Stockholm University) (31). The degree of membrane integration of each inserted segment was quantified as described before (21,22). Samples without CRM were used as controls since proteins translated in their absence would remain unglycosylated.

### Endo H treatment

For Endoglycosidase H (Endo H) treatment, the TNT reaction was mixed with 1 *μ*l of 10× glycoprotein denaturing buffer and dH_2_O to a total volume of 10 *μ*l. After addition of 7 *μ*l of dH_2_O and 2 *μ*l of 10× G5 reaction buffer, samples were incubated for 1 h at 37°C in the presence or absence (mock) of 0.5 *μ*l of Endo Hf enzyme (10^6^ U/ml; NEB, USA).

### Analysis and quantitation

The degree of membrane integration of each TM segment was quantified as described previously (21,22). From the quantified glycosylated products measured ΔG values could be calculated. Fractions of mono- *(f*_*1x*_) and di- *(f*_*2x*_) glycosylated species were quantified in order to calculate an apparent equilibrium constant K_*app*_ for the membrane insertion of the segment. For LepH3 *K*_*app*_ is calculated as 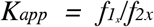 and for LepH2 the fractions of cleaved f_c_ molecules were accounted for and *K*_*app*_ is calculated as *K*_*app*_ = *f*_*2x*_+*f*_*c*_/*f*_*1x*_. *K*_*app*_ can be converted into apparent free energies, measured *ΔG*= -*RTlnK*_*app*_, between the inserted and non-inserted states.

### Cell culture and transfections

HEK293T cells were cultured in DMEM (Life Technologies, USA) supplemented with 10% FBS (Sigma-Aldrich Merck, USA), 100 units/ml penicillin and 100 μg/ml streptomycin (Sigma-Aldrich Merck, USA). The cells were cultured in a humidified 5% CO_2_ incubator at 37°C. For the transfections, 1.5 μg of plasmid DNA was mixed with 1 ml of opti-MEM (Life Technologies, USA) and 5 μl Trans-IT®LT-1 transfection reagent (Mirus, USA) for 20 min. Trypsinized HEK293T cells were resuspended in opti-MEM 10% FBS, and 1 ml of cells of approximate density 10^6^/ml, was added to each transfection mixture before plating on a 3.5 cm diameter dish.

### Cell lysate preparation

Cells were harvested 24 hours after transfection by scraping in 150 μl of Pierce™ RIPA buffer (ThermoFisher Scientific, USA) supplemented with N-ethylmaleimide (Sigma-Aldrich Merck, USA) and 1% cOmplete protease inhibitor (Roche, Germany), sonicated on ice 3 times for 5 seconds and centrifuged for 5 minutes at 15,000 rpm at 4°C. The supernatant was subjected to analysis or stored at −20°C long-term.

### Endoglycosidase H treatment and Western blot on lysates from HEK293T cells

For Endoglycosidase H (Endo H) treatment, 18 μl of protein lysate was mixed with 2 μl of sodium acetate (800 mM, pH 5.7), and 0.75 μl of Endo H (500,000 units/ml; NEB, MA, USA) or dH_2_O (mock sample). The samples were incubated at 37°C for 2 hours, mixed with Werner’s sample buffer and denatured by heating for 5 minutes at 95°C. SDS–PAGE and Western blot analysis with a 1:3333 dilution of a rabbit anti-GFP antibody (Rockland, USA) were carried out. Western blots were developed with SuperSignal™ West Femto Maximum Sensitivity Substrate (ThermoFisher Scientific, USA) and detected with an Azure c600 (Azure Biosystems, USA) imaging system.

### Cell imaging

HEK293T transfected cells cultured on coverslips were fixed 24 hours post-transfection with 4% formaldehyde, washed 3 times with 1x PBS and mounted on glass slides with VECTASHIELD® Antifade Mounting Medium with DAPI (Vector Laboratories, USA). Images were acquired using Zen software connected to an inverted microscope (LSM700; ZEISS, Germany) with 405, and 488 solid-state lasers and 20x objective. For the negative control the zoom out option of ZEN software was used.

## Acknowledgments

We gratefully thank Dr. Rickard Hedman for providing EasyQuant and Prof. Arthur E. Johnson for providing rough microsomes. We also gratefully thank Dr. Hunsang Lee and Dr. Hyun Kim for their advice and suggestions on how to analyze the fluorescent data of the gGFP-assay.

## Author contributions

Planned experiments: (ÅT-R, AA, IMN, AM, RK, PL, SG, MW), performed experiments: (AA, AM, ÅT-R, RK, QL, PL, JP, XR, PX), analyzed data: (ÅT-R, AA, AM, PL, IMN, RK), wrote the paper: (ÅT-R, RK, IMN, GvH, MW, AM, TH).

## FOOTNOTES

This work was supported by grants from the Swedish Cancer Society (130624) to IMN, from the Swedish Foundation for International Cooperation in Research and Higher Education (STINT) (210/083(12) and KU 2003-4674) to IMN, from the Swedish Foundation for Strategic Research (A302:200) to IMN and SSF-Infection Biology 2012(SB12-0026) to IMN and MW, from the Swedish Foundation for Strategic Research (IB13-0026) to TH, and from the Knut and Alice Wallenberg Foundation (2012.0282) and the Swedish Research Council (621-2014-3713) to GvH.

The abbreviations used are: CRM, column-washed rough microsomes from dog pancreas; OST, oligosaccharyl transferase; ER, endoplasmic reticulum; pTM, predicted transmembrane; TM, transmembrane domain; EndoH, endoglycosidase H; SP, signal peptide

---

**Supporting information Figure S1.**
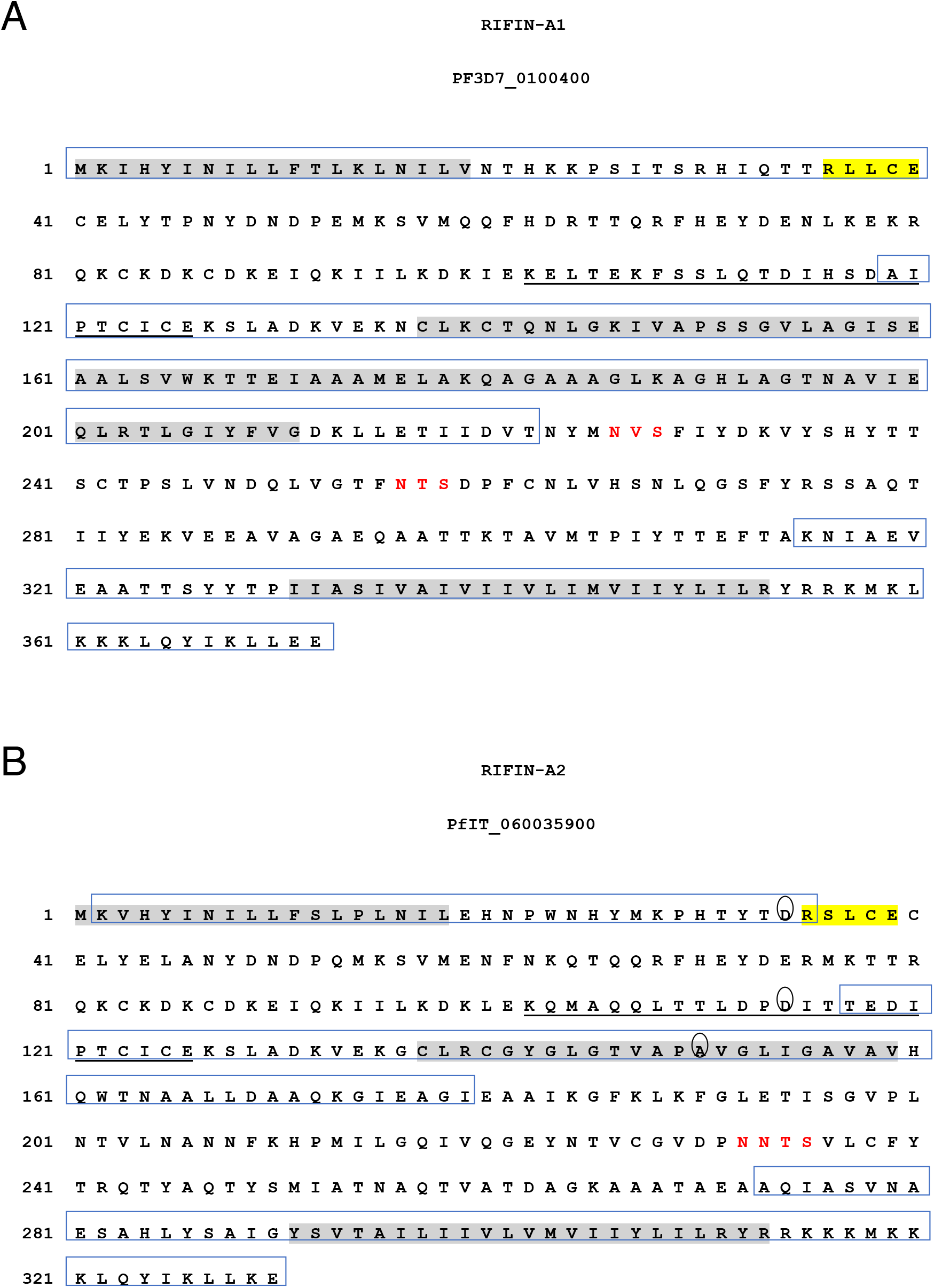
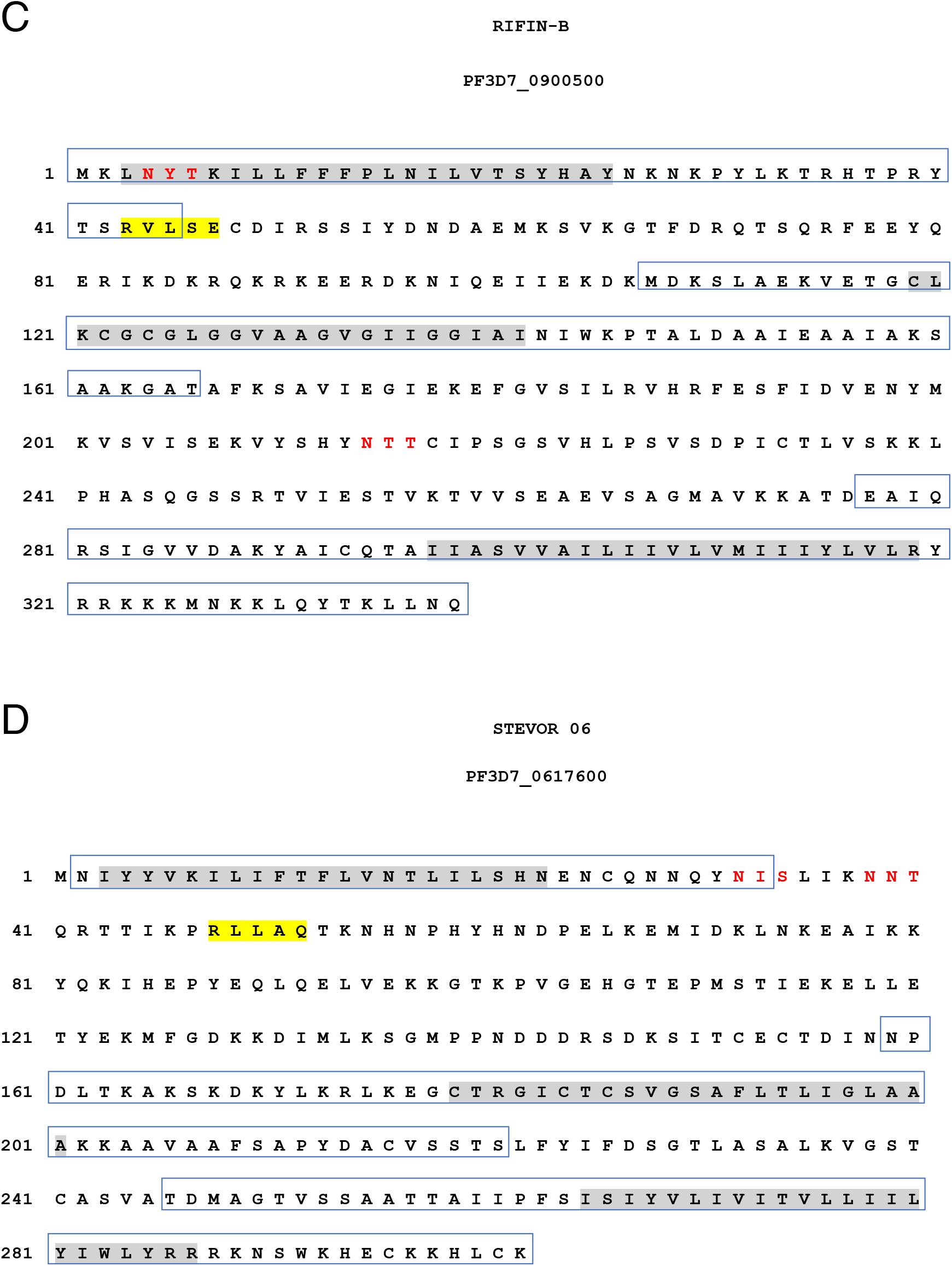
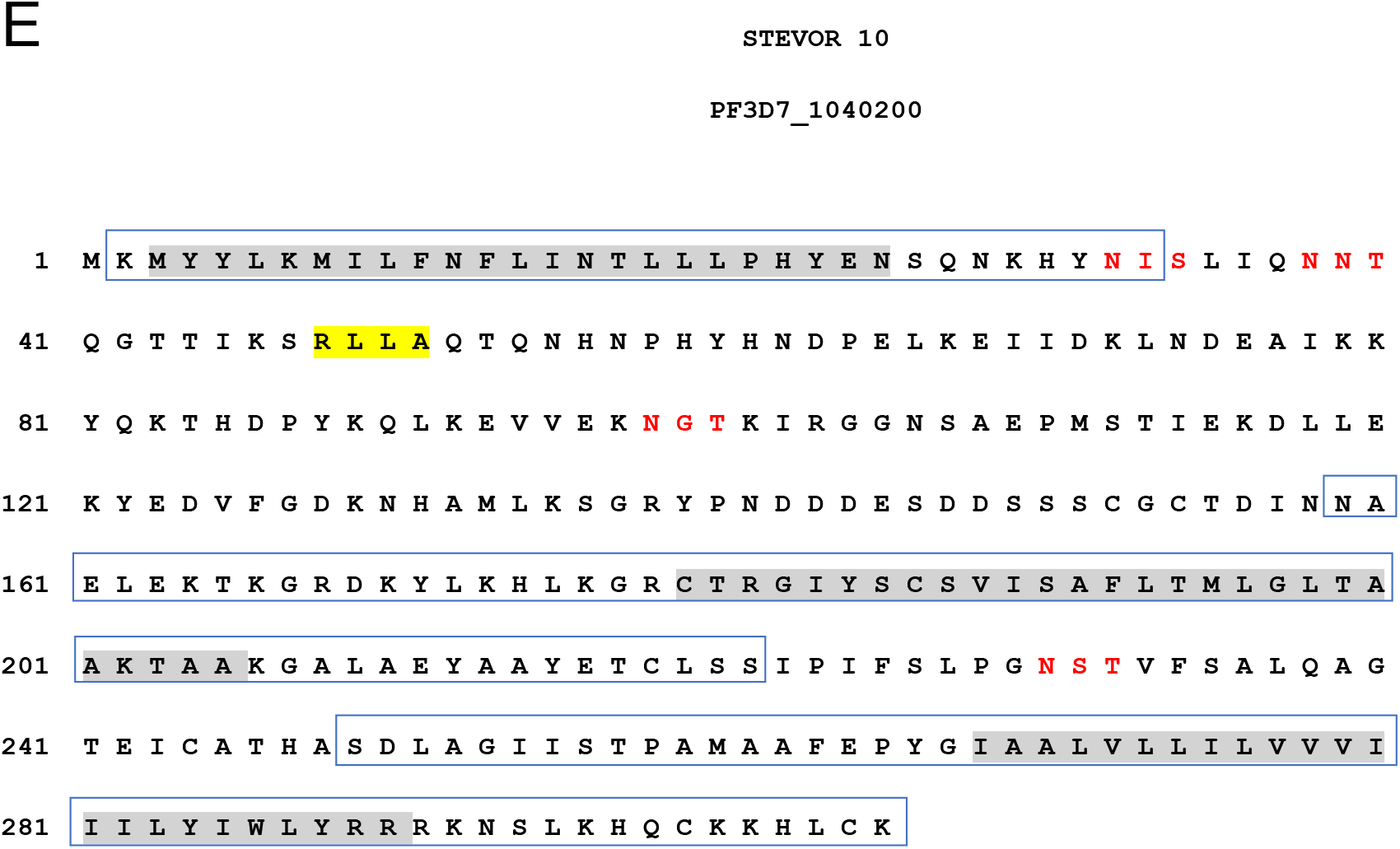
Protein sequences for RIFIN and STEVOR proteins. Predicted TM domains are indicated in gray and the PEXEL site in yellow. Natural glycosylation sites are indicated in red and putative TMs in a box. Underlined sequences in RA1 and 2 represent the indel sequence that is specific for RA.

## References

1. Organisation, W. H. (2018) World Malaria Report 2018. in Geneva: World Health Organization; 2018., Available at http://www.who.int/malaria/publications/world-malaria-report-2018/report/en/

2. Doumbo, O. K., Thera, M. A., Kone, A. K., Raza, A., Tempest, L. J., Lyke, K. E., Plowe, C. V., and Rowe, J. A. (2009) High levels of Plasmodium falciparum rosetting in all clinical forms of severe malaria in African children. Am J Trop Med Hyg 81, 987–993

3. Kaul, D. K., Roth, E. F., Jr., Nagel, R. L., Howard, R. J., and Handunnetti, S. M. (1991) Rosetting of Plasmodium falciparum-infected red blood cells with uninfected red blood cells enhances microvascular obstruction under flow conditions. Blood 78, 812–819

4. Craig, A., and Scherf, A. (2001) Molecules on the surface of the Plasmodium falciparum infected erythrocyte and their role in malaria pathogenesis and immune evasion. Mol Biochem Parasitol 115, 129–143

5. Cheng, Q., Cloonan, N., Fischer, K., Thompson, J., Waine, G., Lanzer, M., and Saul, A. (1998) stevor and rif are Plasmodium falciparum multicopy gene families which potentially encode variant antigens. Mol Biochem Parasitol 97, 161–176

6. Kaur, J., and Hora, R. (2018) ‘2TM proteins’: an antigenically diverse superfamily with variable functions and export pathways. PeerJ 6, e4757

7. Bachmann, A., Scholz, J. A., Janssen, M., Klinkert, M. Q., Tannich, E., Bruchhaus, I., and Petter, M. (2015) A comparative study of the localization and membrane topology of members of the RIFIN, STEVOR and PfMC-2TM protein families in Plasmodium falciparum-infected erythrocytes. Malar J 14, 274

8. Joannin, N., Abhiman, S., Sonnhammer, E. L., and Wahlgren, M. (2008) Sub-grouping and sub-functionalization of the RIFIN multi-copy protein family. BMC Genomics 9, 19

9. Petter, M., Haeggstrom, M., Khattab, A., Fernandez, V., Klinkert, M. Q., and Wahlgren, M. (2007) Variant proteins of the Plasmodium falciparum RIFIN family show distinct subcellular localization and developmental expression patterns. Mol Biochem Parasitol 156, 51–61

10. Niang, M., Bei, A. K., Madnani, K. G., Pelly, S., Dankwa, S., Kanjee, U., Gunalan, K., Amaladoss, A., Yeo, K. P., Bob, N. S., Malleret, B., Duraisingh, M. T., and Preiser, P. R. (2014) STEVOR is a Plasmodium falciparum erythrocyte binding protein that mediates merozoite invasion and rosetting. Cell Host Microbe 16, 81–93

11. Tuteja, R. (2007) Unraveling the components of protein translocation pathway in human malaria parasite Plasmodium falciparum. Archives of biochemistry and biophysics 467, 249–260

12. Tuteja, R., Pradhan, A., and Sharma, S. (2008) Plasmodium falciparum signal peptidase is regulated by phosphorylation and required for intra-erythrocytic growth. Mol Biochem Parasitol 157, 137–147

13. Kelleher, D. J., and Gilmore, R. (2006) An evolving view of the eukaryotic oligosaccharyltransferase. Glycobiology 16, 47R–62R

14. Macedo, C. S., Schwarz, R. T., Todeschini, A. R., Previato, J. O., and Mendonca-Previato, L. (2010) Overlooked post-translational modifications of proteins in Plasmodium falciparum: N- and O-glycosylation -- a review. Mem Inst Oswaldo Cruz 105, 949–956

15. Marti, M., Good, R. T., Rug, M., Knuepfer, E., and Cowman, A. F. (2004) Targeting malaria virulence and remodeling proteins to the host erythrocyte. Science 306, 1930–1933

16. Tarr, S. J., and Osborne, A. R. (2015) Experimental determination of the membrane topology of the Plasmodium protease Plasmepsin V. PLoS One 10, e0121786

17. Russo, I., Babbitt, S., Muralidharan, V., Butler, T., Oksman, A., and Goldberg, D. E. (2010) Plasmepsin V licenses Plasmodium proteins for export into the host erythrocyte. Nature 463, 632–636

18. Kirk, K., and Lehane, A. M. (2014) Membrane transport in the malaria parasite and its host erythrocyte. Biochem J 457, 1–18

19. Elsworth, B., Crabb, B. S., and Gilson, P. R. (2014) Protein export in malaria parasites: an update. Cell Microbiol 16, 355–363

20. Goel, S., Palmkvist, M., Moll, K., Joannin, N., Lara, P., Akhouri, R. R., Moradi, N., Ojemalm, K., Westman, M., Angeletti, D., Kjellin, H., Lehtio, J., Blixt, O., Idestrom, L., Gahmberg, C. G., Storry, J. R., Hult, A. K., Olsson, M. L., von Heijne, G., Nilsson, I., and Wahlgren, M. (2015) RIFINs are adhesins implicated in severe Plasmodium falciparum malaria. Nat Med 21, 314–317

21. Hessa, T., Kim, H., Bihlmaier, K., Lundin, C., Boekel, J., Andersson, H., Nilsson, I., White, S. H., and von Heijne, G. (2005) Recognition of transmembrane helices by the endoplasmic reticulum translocon. Nature 433, 377–381

22. Lundin, C., Kim, H., Nilsson, I., White, S. H., and von Heijne, G. (2008) Molecular code for protein insertion in the endoplasmic reticulum membrane is similar for N(in)-C(out) and N(out)-C(in) transmembrane helices. Proc Natl Acad Sci U S A 105, 15702–15707

23. Kozak, M. (1989) Context effects and inefficient initiation at non-AUG codons in eucaryotic cell-free translation systems. Mol Cell Biol 9, 5073–5080

24. Lundin, C., Nordstrom, R., Wagner, K., Windpassinger, C., Andersson, H., von Heijne, G., and Nilsson, I. (2006) Membrane topology of the human seipin protein. FEBS Lett 580, 2281–2284

25. Johansson, M., Nilsson, I., and von Heijne, G. (1993) Positively charged amino acids placed next to a signal sequence block protein translocation more efficiently in Escherichia coli than in mammalian microsomes. Mol Gen Genet 239, 251–256

26. Hessa, T., Meindl-Beinker, N. M., Bernsel, A., Kim, H., Sato, Y., Lerch-Bader, M., Nilsson, I., White, S. H., and von Heijne, G. (2007) Molecular code for transmembrane-helix recognition by the Sec61 translocon. Nature 450, 1026–1030

27. Arthur, L., Pavlovic-Djuranovic, S., Smith-Koutmou, K., Green, R., Szczesny, P., and Djuranovic, S. (2015) Translational control by lysine-encoding A-rich sequences. Sci Adv 1

28. Lee, H., Lara, P., Ostuni, A., Presto, J., Johansson, J., Nilsson, I., and Kim, H. (2014) Live-cell topology assessment of URG7, MRP6 and SP-C using glycosylatable green fluorescent protein in mammalian cells. Biochem Biophys Res Commun

29. Walter, P., and Blobel, G. (1983) Preparation of microsomal membranes for cotranslational protein translocation. Methods Enzymol 96, 84–93

30. Cuviello, F., Tellgren-Roth, A., Lara, P., Ruud Selin, F., Monne, M., Bisaccia, F., Nilsson, I., and Ostuni, A. (2015) Membrane insertion and topology of the amino-terminal domain TMD0 of multidrug-resistance associated protein 6 (MRP6). FEBS Lett 589, 3921–3928

31. Ismail, N., Hedman, R., Schiller, N., and von Heijne, G. (2012) A biphasic pulling force acts on transmembrane helices during translocon-mediated membrane integration. Nat Struct Mol Biol 19, 1018–1022

32. Karamyshev, A. L., Kelleher, D. J., Gilmore, R., Johnson, A. E., von Heijne, G., and Nilsson, I. (2005) Mapping the interaction of the STT3 subunit of the oligosaccharyl transferase complex with nascent polypeptide chains. J Biol Chem 280, 40489–40493

33. Seppala, S., Slusky, J. S., Lloris-Garcera, P., Rapp, M., and von Heijne, G. (2010) Control of membrane protein topology by a single C-terminal residue. Science 328, 1698–1700

34. von Heijne, G. (1986) The distribution of positively charged residues in bacterial inner membrane proteins correlates with the trans-membrane topology. EMBO J 5, 3021–3027

35. Higy, M., Junne, T., and Spiess, M. (2004) Topogenesis of membrane proteins at the endoplasmic reticulum. Biochemistry 43, 12716–12722

36. Goder, V., and Spiess, M. (2003) Molecular mechanism of signal sequence orientation in the endoplasmic reticulum. Embo J 22, 3645–3653

37. Gafvelin, G., Sakaguchi, M., Andersson, H., and von Heijne, G. (1997) Topological rules for membrane protein assembly in eukaryotic cells. J Biol Chem 272, 6119–6127

38. Gafvelin, G., and von Heijne, G. (1994) Topological “frustration” in multi-spanning *E. coli* inner membrane proteins. Cell 77, 401–412

39. Przyborski, J. M., and Lanzer, M. (2005) Protein transport and trafficking in Plasmodium falciparum-infected erythrocytes. Parasitology 130, 373–388

40. Przyborski, J. M., Miller, S. K., Pfahler, J. M., Henrich, P. P., Rohrbach, P., Crabb, B. S., and Lanzer, M. (2005) Trafficking of STEVOR to the Maurer’s clefts in Plasmodium falciparum-infected erythrocytes. EMBO J 24, 2306–2317

41. Sleebs, B. E., Lopaticki, S., Marapana, D. S., O’Neill, M. T., Rajasekaran, P., Gazdik, M., Gunther, S., Whitehead, L. W., Lowes, K. N., Barfod, L., Hviid, L., Shaw, P. J., Hodder, A. N., Smith, B. J., Cowman, A. F., and Boddey, J. A. (2014) Inhibition of Plasmepsin V activity demonstrates its essential role in protein export, PfEMP1 display, and survival of malaria parasites. PLoS Biol 12, e1001897

